# Online modification of goal-directed control in human reaching movements

**DOI:** 10.1101/2020.09.03.280784

**Authors:** Antoine De Comite, Frédéric Crevecoeur, Philippe Lefèvre

**Author notes:** ADC, FC and PL designed the study, analysed the data and drafted the manuscript, ADC collected the data. Corresponding author : Philippe Lefèvre, 4 Avenue Georges Lemaître, 1348 Louvain-la-Neuve, Belgium, Mail.

## Abstract

Humans are able to perform very sophisticated reaching movements in a myriad of contexts based on flexible control strategies influenced by the task goal and environmental constraints such as obstacles. However, it remains unknown whether these control strategies can be adjusted online. The objective of this study was to determine whether the factors which determine control strategies during planning also modify the execution of an ongoing movement following sudden changes in task demand. More precisely, we investigated whether, and at which latency, feedback responses to perturbation loads followed the change in the structure of the goal target or environment. We changed the target width (square or rectangle) to alter the task redundancy, or the presence of obstacles to induce different constraints on the reach path, and assessed based on surface recordings when the change in visual display altered the feedback response to mechanical perturbations. Task-related EMG responses were detected within 150 ms of a change in target shape. Considering visuomotor delays of ∼ 100 ms, these results suggest that it takes 50 ms to change control policy within a trial. An additional 30 ms delay was observed when the change in context involved sudden appearance or disappearance of obstacles. Overall, our results demonstrate that the control policy within a reaching movement is not static: contextual factors which influence movement planning also influence movement execution at surprisingly short latencies. Moreover, the additional 30 ms associated with obstacles suggest that these two types of changes may be mediated via distinct processes.

**New & Noteworthy:** The present work demonstrates that the control strategies used to perform reaching movements are adjusted online when the structure of the target or the presence of obstacles are altered during movements. Thus, the properties of goal-directed reaching control are not simply selected during the planning stage of a movement prior to execution. Rather, they are dynamically and rapidly adjusted online, within ∼150ms, according to changes in environment.

## Introduction

Humans perform dozens of reaching movements every day in a broad range of contexts. It has been demonstrated that the properties of these movements depend on various parameters such as the size and orientation of a target and the presence of obstacles (Chapman and Goodale 2008; Glazebrook 2018; Howard and Tipper 1997; Jackson et al. 1995; Knill et al. 2011; Lacquaniti et al. 1986; Nashed et al. 2012). Indeed, these factors influence planning strategies as reflected in the ability of both visual and mechanical perturbations to affect the kinematics of unperturbed movements and feedback control strategies. For example, when reaching for a large rectangular target, participants do not correct deviations aligned with the larger dimension of that target as they do when reaching for a small square (Knill et al. 2011; Nashed et al. 2012). Similarly, participants modify their reaching trajectories to navigate around obstacles dependent on inertial factor (Sabes et al. 2018) and following perturbations (Nashed et al. 2012). Both sets of results provide experimental evidence that control strategies used by humans depend on task goals and environmental context. These results also demonstrate that a continuous stream of sensory information regarding a task and its environment is translated into the neural feedback controller (Crevecoeur et al. 2019; Izawa and Shadmehr 2008; Scott 2016).

Many previously conducted experiments have been motivated by the theory of optimal feedback control. This theory suggests that the control strategy used to perform movement is tuned to the task goal and to its environmental context. In this framework, task constraints and knowledge of limb dynamics are used to derive an optimal goal-dependent control strategy that defines the feedback gains applied during movement (Liu and Todorov 2007; Scott 2004; Todorov and Jordan 2002). It is proposed that the goal-dependent control policy produces both the voluntary motor commands to the target as well as corrective feedback responses to external perturbations. Optimal feedback control can reproduce many of the experimental features detailed above by considering that different cost-functions correspond to different tasks. It is important to realize that in previous studies, the cost-functions used to characterize task-dependent control policies were typically varied across conditions, including for instance different targets in a tracking task or different target shapes (Gallivan et al. 2016b; Lowrey et al. 2017; Nashed et al. 2012, 2014). These previous studies have not considered change in cost function corresponding to change in task demands occurring during a movement. However, under natural conditions, it is conceivable that control policies and cost parameters vary over time. For example, in field sports the obstacles (e.g., opponents) move quite a bit. To date, it remains unknown how quickly the nervous system can update its control policy following a change in cost-function.

Dynamic updates in cost-function, followed by adjustments in control policy, have been described in theory in the framework of model predictive control ((Lee 2011) for review). The basic premise of this theory is that changes are addressed with successive re-computations of control policy. This framework was used by Dimitriou and colleagues (Dimitriou et al. 2013) to interpret how feedback gains are updated in a reaching task when both the goal target and a hand-aligned cursor jumped during movements. These authors observed changes in feedback gains when the target jumped during movement suggesting an online re-computation of control policy evoked in response to jumping of the target. It remained unclear how quickly the control strategy was re-computed, which was the main purpose of our study.

To address this issue, a specific paradigm was needed because a possible shortcoming of the approach based on target jump is that it does not necessarily imply a re-computation of feedback gains (Li et al. 2018). Instead, feedback responses in this case could result from state feedback control associated with long (infinite) time horizon (Qian et al. 2013). Therefore, in order to test how quickly the nervous system can update its control policy, a perturbation which alters not only the state of the goal (i.e. the target location), but also its structure is needed. In this manner, feedback responses to perturbations do not simply follow from a state-feedback correction but also involve an adjustment of the control policy.

To achieve this goal, we leveraged the impact of a change in target structure on reach control to investigate whether an online modification of goal evokes an adjustment of feedback gains. Specifically, the target was switched during movement from a small square to a large rectangle. This switch represented a change from constraints imposed on two coordinates of the endpoint to redundancy along the main axis, respectively. In this context, two alternative hypotheses exist: if the control policy remains fixed within each movement, we expect that a change in target shape will have no impact on feedback responses to perturbation loads, and no changes in movement kinematics should be detected. In contrast, if a change in task demand evokes changes in the control policy online, then we expect that the feedback responses to the mechanical loads and subsequent kinematics adjust to the new target. Consistent with the latter hypothesis, we found that participants produced corrective responses adjusted to the change in target shape as well as to changes in environmental context (e.g., the presence of obstacles). Strikingly, evoked changes in muscle activity were observed in as little as 150 ms following the change in target shape, while we measured a slightly longer time needed to produce an adjusted response with obstacles. This result demonstrates that control policy during reaching can be updated rapidly and during movement execution.

## Methods

### Participants

Three groups of participants were enrolled in this study. The first group performed experiment 1 and included 14 right-handed participants (5 females, 9 males) ranging in age from 18 to 29 years. The second group performed experiment 2 and included 17 right-handed participants (7 females, 10 males) ranging in age from 19 to 25 years. The last group performed the control experiment and included 8 right-handed participants (4 females, 4 males) ranging in age from 19 to 26 years. All sets of participants were naïve to the purpose of the experiments and had no known neurological disorders. The local ethics committee of the Université catholique de Louvain approved the experimental procedures of this study.

### Setup

Participants sat comfortably on an adjustable chair in front of a KINARM end-point robotic device (BKIN Technologies, Kingston, ON, Canada). The participants were to grasp the handle of the right robotic arm with their right hand. The robotic arm allowed movements in the horizontal plane. Direct vision of both the participant’s hand and the robotic arm was blocked. Participants were seated such that their elbow, pointing down to the ground, formed an angle of approximately 90° and their forehead was placed on a soft cushion attached to the frame of the setup. A virtual reality display was placed above the handle to allow the participants to interact with virtual targets and obstacles. A white dot with a radius of 0.5 cm corresponded to the position of the participant’s hand throughout the experiment.

### Experiment 1

In the first experiment (see Figure 1, left panel), participants (n = 14) were instructed to perform reaching movements to a visual target that was either a small square (2.5 cm x 2.5 cm) or a large rectangle (30 cm x 2.5 cm) located 25 cm in the y-direction from the square home target (2.5 cm x 2.5 cm). The main axis of the rectangle was aligned with the x-axis and was orthogonal to the straight-line path from the starting point to the target goal. Briefly, participants had to put a hand-aligned cursor on the home target (displayed as a red square). When the home target was reached, the square turned green. After a random time delay (anywhere from 2–4 s), the goal target was projected and participants could begin movement whenever they wanted. Thus, there were no constraints on reaction time. Following exit of the home target, participants had to complete their movements between 350 ms and 600 ms from the start of movement in order to successfully complete the trial. The trial was successfully completed if participants reached the goal target within the prescribed window and were able to stabilize the cursor in it for 500 ms. The goal target turned green at the end of a successful trial. For trials in which the timing window was not respected, the goal target turned blue or red to indicate that the movements performed were too fast or too slow, respectively.

**Figure 1.**
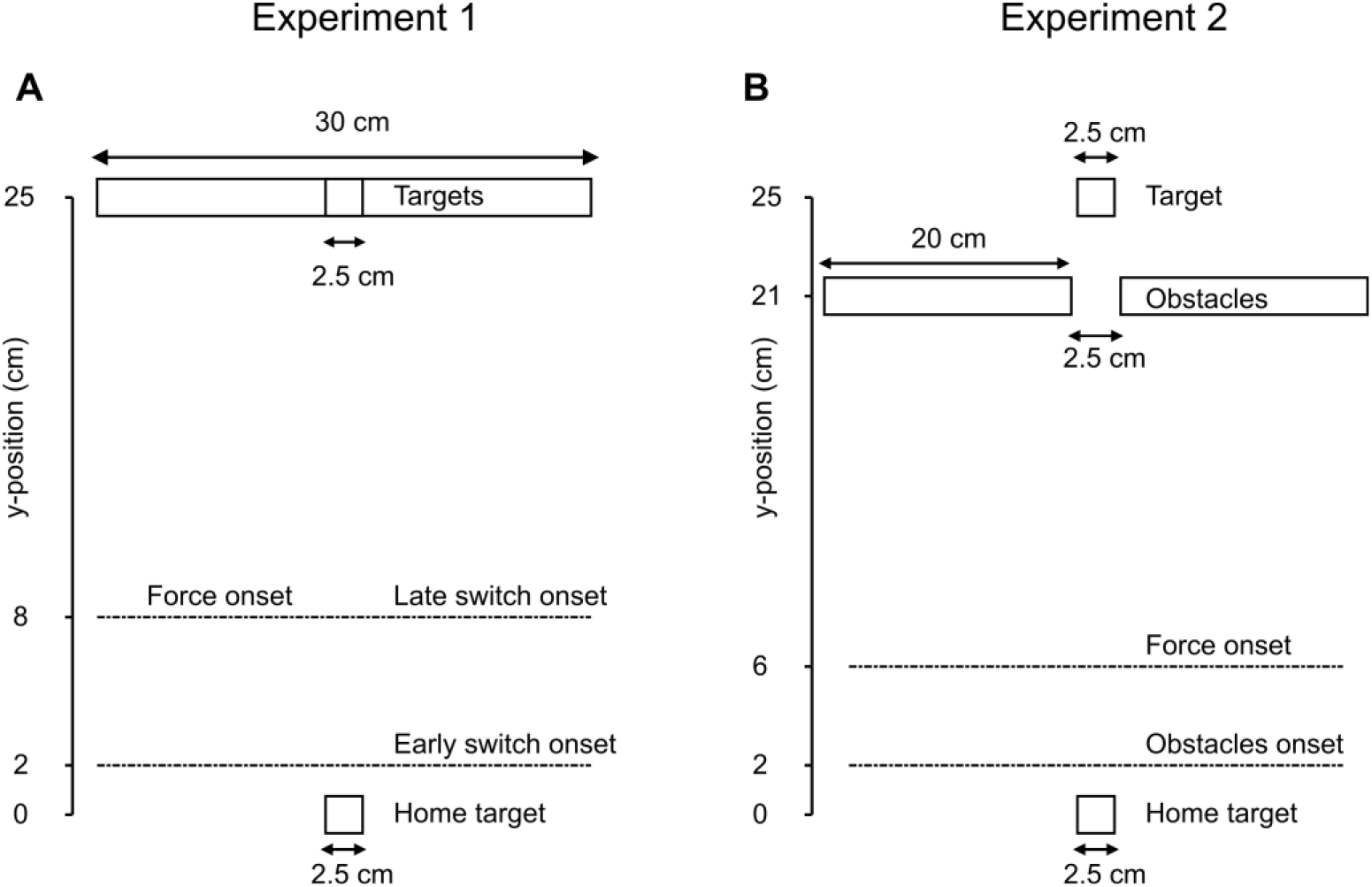
Experimental paradigm. **A**. Schematic of Experiment 1, participants were asked to perform reaching movements from the home target to the goal target (square or rectangle). The dashed lines represent the y-position at which an early switch in target shape, a late switch in target shape, and onset of a force (perturbation load) were introduced. To succeed the trial, participants had to reach and stabilise in the goal target in the prescribed time window. **B**. Schematic of Experiment 2, participants were asked to perform reaching movements from the home target to the goal target while avoiding obstacles if present. The dashed lines indicate the y-position at which obstacles or onset of a force (perturbation load) were introduced. To succeed the trial, participants had to reach the goal target in the prescribed time window without hitting the obstacles.

During the movement, two types of perturbations could occur. The first was a mechanical perturbation load consisting of a lateral step force applied by the robot to the hand of the participant. The magnitude of this force was ± 9N, with a 10-ms linear build-up aligned with the x-axis. This force was triggered when the hand-aligned cursor crossed a virtual line that was aligned with the x-axis and was located 8 cm from the center of the home target (see Figure 1 left panel). The step force was switched off at the end of the trial. The second type of perturbation applied was a visual perturbation where the shape of the target was instantaneously changed from a small square to a large rectangle, or vice versa. This switch in target shape was also triggered based on a position threshold, positioned either 2 cm (early target switch) or 8 cm (late target switch) from the center of the home target. Both perturbed and unperturbed trials were randomly interleaved such that participants could not predict the occurrence of either visual or mechanical perturbations.

Participants started with a training block of 20 trials in order to become familiar with the task and the force intensity of the perturbation loads. After completing the training block, six blocks of 72 trials were conducted. Each 72-trial block included: 24 trials involving no force or target switch, 16 trials with a mechanical perturbation only, and 32 trials with both visual and mechanical perturbations. Thus, participants completed 432 trials, amongst which 24 trials of each perturbation combination were included [e.g., direction of the mechanical perturbation changed (e.g., rightward versus leftward) and timing of the target changed (e.g., early versus late)]. Participants were verbally encouraged to exploit the width of the target when it was a rectangle. To motivate the participants, a score corresponding to their number of successful trials was projected next to the goal target.

### Experiment 2

The second experiment was a variant of the first one and was designed to assess reproducibility in a distinct context. Thus, instead of changing the structure of the goal, obstacles were introduced (see Figure 1, right panel). Briefly, participants had to perform reaching movements towards a square target (2.5 cm x 2.5 cm) which was positioned similarly to Experiment 1. However, two virtual obstacles (20 cm x 2.5 cm each) were located 2.5 cm apart on either side of the target reach path, with their main axes aligned with the x-axis and orthogonal to the main reaching direction. When participants hit these obstacles, a reaction force pushed their hand back to provide haptic feedback and they lost a point. Similar to the first experiment, participants had to reach the home target to start the trial. After a random time delay, the goal target appeared as a red square and they could start their movement whenever they wanted. Again, two types of perturbations were applied during a movement. The first perturbation was the same mechanical perturbation used in Experiment 1, and it was triggered 6 cm from the center of the home target and was aligned with the x-direction (positive or negative). We observed in some pilot testing that in this context, the success rate was very low if the mechanical perturbation was triggered at 8 cm from the center of the home target. For this reason, we decided to trigger this perturbation at 6 cm. The second type of perturbation involved the appearance or disappearance of obstacles. These perturbations were triggered when the hand of a participant crossed a virtual line which was aligned with the x-axis and located 2 cm from the center of the home target. All trials were randomly interleaved. Following a similar procedure as Experiment 1, participants first performed a training block of 20 trials involving both force and obstacles changes. After completing this training block, participants performed six blocks of 72 trials. Each 72-trial block included: 32 unperturbed trials, 16 trials with a mechanical perturbation but no obstacles, and 24 trials which contained both mechanical perturbations (e.g., leftward or rightward) and visual perturbations with an appearance or disappearance of obstacles. Participants performed a total of 432 trials, including 24 trials which included each perturbation combination (e.g., direction of the mechanical perturbation changed (e.g., rightward versus leftward) and appearance or disappearance of obstacles). Also similar to Experiment 1, the number of successfully completed trials was projected next to the initial target to help motivate the participants.

### Experiment 3

The third experiment was a control experiment designed to verify that the change in behavior observed in Experiment 1 could be elicited without mechanical perturbations and that therefore it was not a consequence of the perturbation load applied to the arm specific to feedback control. Thus, this control experiment was a variant of Experiment 1, in which there was no mechanical perturbation. Similar to Experiment 1, participants (N=8) had to reach for either a small square or a large rectangular target located 20 cm in the y-direction from the home target. The timing constraints were the same as in Experiment 1: between 350 and 600 ms to complete movement once the hand-aligned cursor exited the home target. During movement, changes in target structure as in Experiment 1 could occur: the shape of the goal target could instantaneously change from a square to a rectangle or vice-versa. This change was triggered when participant’s hand crossed a line located at 5 cm from the home target. Movements with or without target switch were randomly interleaved and the visual perturbations occurred in one of three trials. Participants first performed a training block of 20 trials without target switch, followed by three blocks of 60 trials consisting of 40 trials without online target switch, 10 trials with online switch from square to rectangle, and 10 trials with switches from rectangle to square per block.

In all experiments, mechanical and visual perturbations were triggered based on a position threshold in an attempt to reduce variability in joint configuration and electromyography data (EMG). Both kinematics and EMG data were aligned according to the onset of mechanical perturbation prior to data analysis. Since the early switch condition and the appearance/disappearance of obstacles were not triggered at the same position threshold as the mechanical perturbation, there was some variability in the onset of these perturbations. Therefore, the early target switch was triggered 82.45 ± 5.11 ms before the force onset. The late switch condition was triggered at the same time as the mechanical load. As a result, onset was fixed with respect to the onset of the mechanical perturbation. However, since visual processing of the setup was necessary, there was a 40 ms delay between triggering of the visual change (namely the change in target shape or the appearance/disappearance of obstacles) and the visual change on the virtual reality display due to hardware limitations. In this paper, this 40 ms delay was taken into account in all the analyses, which means that the onsets of visual changes (both change in target shape and in environmental context) reported have been delayed by 40 ms with respect to their respective triggers.

### Data recording

Raw kinematics data were sampled at 1 kHz and low-pass filtered with a 4^th^ order double-pass Butterworth filter (cut-off frequency, 20 Hz). Hand velocities along the x- and y-axes were computed from numerical differentiation of the position data by using a 4^th^ order centered finite difference. The raw EMG data were band-pass filtered with a 4^th^ order double-pass Butterworth filter with a band pass between 20 Hz and 250 Hz. EMG data were normalized for each participant to the average activity collected when participants maintained postural control at the home target against a constant force of 9 N. The data from the pectoralis major was normalized to the average activity in the same muscle while maintaining postural control against a rightward 9 N force whereas data from deltoid posterior was normalized to the average activity recorded while maintaining postural control against a leftward 9 N force. We measured the muscle activity from 500 ms to 2000 ms after force onset in the calibration trials. Analysis of EMG responses was performed after aligning the trials to the onset of force (see above). Data processing and parameter extractions were performed by using Matlab 2016b software. Mixed model analyses were performed by using R (version 3.4.1) and the package nlme (Pinheiro, Bates, DebRoy, Sarkar & CoreTeam). The type of trial was considered as a fixed effect while participants were considered as random effects. These models were fitted by maximizing the log-likelihood and significance was considered when the p-value of the t-test on the fixed effect term was smaller than 0.05 for the first significant time bin and 0.005 for the following.

In the first two experiments, surface EMG electrodes (Bagnoli Surface EMG Sensor, Delsys Inc., Natick, MA, USA) were used to record muscle activity during movements. Based on previous studies [(Lowrey C.R., 2017) (Nashed J.Y., 2012)], we concentrated on the pectoralis major (PM) and posterior deltoid (PD). These muscles are stretched by the application of the lateral forces applied in our experiments, and therefore, they are largely recruited for the feedback response. The PM was calibrated with a rightward force and the PD was calibrated with a leftward force. This calibration procedure was applied after the second and fourth blocks. The skin of each participant was cleaned and abraded with cotton wool and alcohol before electrodes were applied. Conduction gel was subsequently spread on the electrodes to improve signal quality. EMG data were sampled at a frequency of 1 kHz and were amplified by a factor of 1 000. In both experiments, a reference electrode was attached to the right ankle of the participant. No EMG data was collected in the control experiment.

During the first two experiments, we also collected activity data for other muscles for exploratory analyses. These additional muscles included the triceps lateralis and brachioradialis during the first experiment, and the anterior deltoid, teres major, infraspinatus, and biceps during the second experiment. For these other muscles, the patterns of response were not stereotyped across the participants, likely because the joint configuration was not precisely controlled. Thus, the results described below concentrate on the two muscles for which a clear stretch response could be measured, as well as correlates of changes in behavior as measured with movement kinematics.

## Statistical analysis

Median values of end-point distributions were analysed for statistically significant differences by using Wilcoxon sign ranked test. Distributions obtained under early and late conditions were compared with distributions obtained without changes in target shape for the same force. Two-sample Kolmogorov-Smirnov tests were applied to compare the cumulative density functions of end-point distributions for the participants’ hands at the end of the trial to identify statistically significant differences. To compare hand paths, the Peacock test was applied (Peacock J.A., 1983). This multidimensional expansion of the two-sample Kolmogorov-Smirnov test was used to determine whether two given multidimensional distributions significantly differed. The null hypothesis of this test is that the two sets of multidimensional data points come from the same distribution. This test is similar to the one used for univariate distributions, although it is performed in the two dimensions of the workspace and the largest difference is used to assess whether samples differ. To perform this test, we implemented the algorithm developed by Xiao (Xiao 2017).

To assess feedback responses for conditions with or without a change in target, normalized EMG data were used. For each participant, their EMG data was averaged across trials and across time with ten non-overlaying bins of 25 ms width. The width of the bins has been determined based on prior testing in which we tested bins of 15 ms, 25 ms and 40 ms width. We observed qualitatively very similar results for the three tested bin sizes but noticed that small bins are more sensitive to noise whereas wide bins act as a low pass filter that could either over or underestimate onset times.

Therefore, we selected a bin width of 25 ms that gave us a good trade-off between sensitivity and accuracy. Moreover, this width corresponds to the bins selected for similar analyses in previous studies (Kurtzer et al. 2008; Nashed et al. 2012). The first and last 25 ms bins started 50 ms and 275 ms after the onset of force, respectively. Next, we applied a mixed linear model to the time average of the EMG data in each of these bins (see equation 1). The trial condition, *x*_*i*_, was the fixed effect factor (slope) and participants, *z*_*j*_, were the random effect factor (offset). We assessed the difference between conditions by looking at the significance of the fixed effect parameter *β*_1_. The random effect parameter was introduced to integrate the inter-subject variability in EMG activation.

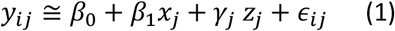

For all of the statistical tests performed, a significance level of p = 0.005 was used, as previously proposed (Benjamin and Berger 2018). To reduce the rate of false negative results, p-values between 0.05 and 0.005 were considered to be indicative of a significant trend. A threshold value of 0.05 was used to determine the onset of differences in the sliding analysis on the basis that bins exhibiting significant differences at this level were followed by bins exhibiting highly significant differences (p < 0.005). Effect size for binned EMG analyses were quantified using Cohen’s d defined as the standardized mean difference between two groups of independent observations (Lakens 2013).

## Results

### Experiment 1

To investigate whether a change in the shape of the goal target during a reaching movement would elicit a change in behavior, participants were instructed to perform movements to reach a target that was either a small square or a large rectangle (see Methods). During these movements, a lateral force could be applied to the hand of participants in order to enhance the effect of a change in target shape on control policy online. The presence and latency of changes in control policy were assessed according to movement kinematics and evoked EMG responses of the muscles stretched by the perturbation loads.

### Kinematics

During the unperturbed trials, movement kinematics depended on the shape of the goal target, consistent with previously reported results (Lowrey et al. 2017; Nashed et al. 2012). For example, the end-point distribution of hand positions along the x-axis (Figure 2C) was wider for the rectangle target (0.84 ± 2.05 cm) than for the square target (−0.11 ± 0.74 cm) (Kolmogorov-Smirnov test, p < 0.005, K = 0.3780). This difference was observed for all of the participants, and trial-by-trial differences were observed for 9 of the 14 participants. Traces of unperturbed trials for a representative participant are shown in Figure 2A. For all of the participants, the reaching time for the small square target (median 422.3 ms) was also longer than that for the large rectangle target (median 389ms) (Wilcoxon signed-rank test, z = 9.64, rank sum = 9.89 × 10^5^, p < 0.001). Trial-by-trial differences in reaching time were observed for 13 of the 14 participants.

**Figure 2.**
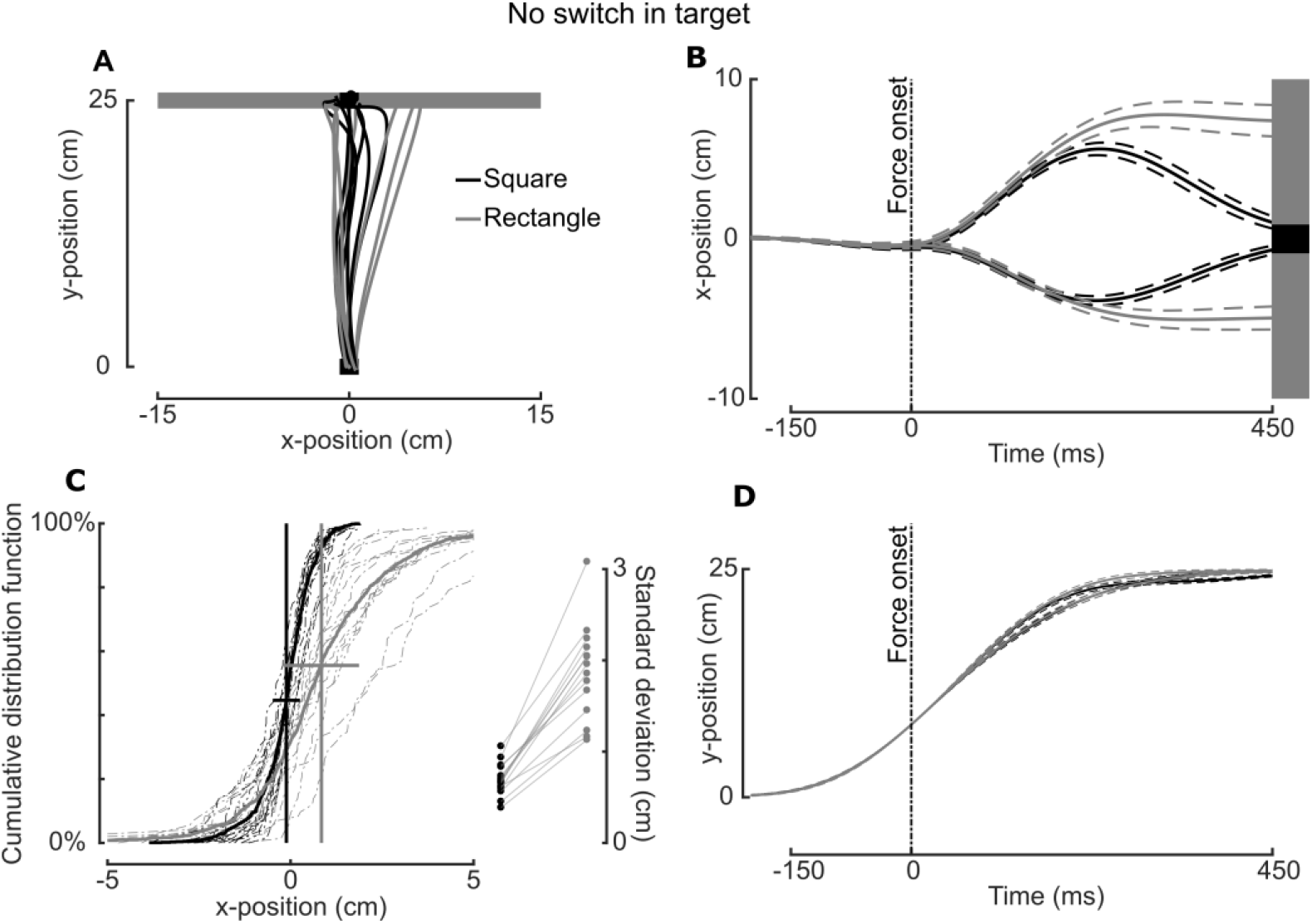
**A**. Hand paths of a representative participant toward a square (black) or rectangle (grey) target without any switch in target shape or perturbation load. **B**. Group mean (solid line) and SEM (dashed line) traces of the hand paths along the x-axis for participants reaching for a square (black) or a rectangle (grey) with perturbation loads applied. The onset of force is indicated with a vertical dashed line. **C**. Left: Cumulative distribution functions of hand end-points for the participants reaching to a square target (black) or a rectangular target (grey) without application of a perturbation load. Solid lines represent mean distribution values, while dotted lines represent individual distribution values. Right: Standard deviations of the individual end-point distributions of participants’ hands reaching to a square (black dots) or rectangle (grey dots) target. **D**. Group mean and SEM y-position values of participants’ hand path when reaching for a square (black) versus rectangle (grey) target with perturbation loads applied. The onset of force is indicated with a vertical dashed line.

Similarly, when perturbation loads were applied without a change in target shape, kinematics of the movement were found to depend on the shape of the target (Figure 2B). For example, the mean x-position of the participants’ hands at the end of the trials involving rightward perturbation loads for the square target was 1.04 ± 1.58 cm (median ± standard deviation). For the rectangle target, the mean x-position was 7.41 ± 4.05 cm (Kolmogorov-Smirnov test, p < 0.005, K = 0.5089). This difference was observed across all of the trials for the participants (Figure 2C). When leftward perturbation loads were applied, a similar difference in the x-positions of the participants’ hands was observed at the end of the trials (Figure 2B). However, no differences in hand position were observed along the y-axis (Figure 2D).

Based on the observation that control strategies used during movement depend on the shape of the goal target, we next wanted to examine whether similar adjustments would be evoked online when the goal target suddenly changed during movement. Therefore, a change in the shape of the target goal from a square to a rectangle, or vice versa, was introduced just after leaving the home target (early switch) or just after the application of a mechanical perturbation load (late switch). Distinct changes in the mean end-point coordinate and distribution were observed.

When an early target switch from square to rectangle was made with rightward perturbation (red in Figure 3, A and B), the mean deviation in hand position along the x-axis was larger than that observed for the late switch condition (6.01 ± 3.54 cm vs. 2.65 ± 2.68 cm, respectively). A comparison with the end-point distribution of the non-switching condition also highlighted different distributions (Kolmogorov-Smirnov test, p < 0.005, K = 0.3155, red versus black in Figure 3A).

**Figure 3.**
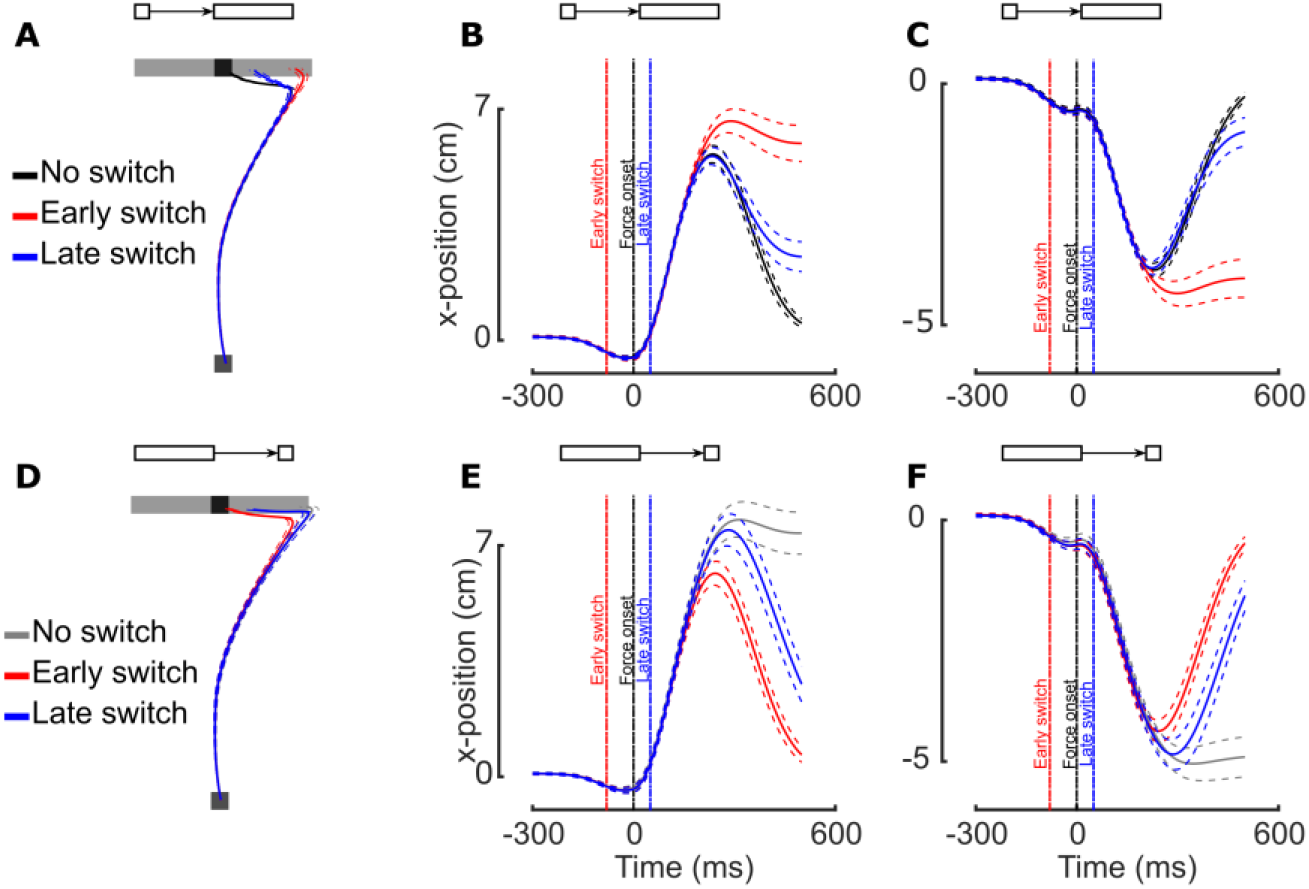
**A-F**. Mean (solid line) and SEM (dashed line) traces of participants’ hand paths (**A**,**D**) or x-positions with respect to time (**B**,**C**,**E**,**F**) in the absence (**A**,**D**) or presence of a rightward (**B**,**E**) or leftward (**C**,**F**) perturbation load (indicated as a vertical dashed line) while reaching for a square (**A**, black), a rectangle (**D**, grey), a square switched to a rectangle (**B**,**C**, black), or a rectangle switched to a square (**E**,**F**, grey). The red and blue traces represent the early and late switch conditions, respectively.

When a late target switch from square to rectangle was made with rightward perturbation (in blue in Figure 3, A and B), a larger deviation in hand position along the x-axis was observed than when a target switch did not occur (in black in Figure 3, A and B). The mean distribution values for the trials with a late target switch and without a target switch were 2.65 ± 2.68 cm and 1.04 ± 1.58 cm, respectively. These two end-point distributions differed across all of the participants (Kolmogorov-Smirnov test, p < 0.005, K = 0.6905, blue versus black in Figure 3B).

Differences in hand deviation along the x-axis were also observed for the switches from rectangle to square targets. Deviations of the participants’ hand positions along the x-axis for both early and late switching conditions were smaller than the deviation for the non-switching condition (see Figure 3E for results obtained with rightward perturbation).

The final position of the participants’ hands also exhibited dependency on when the target was switched. For example, when we compared the final positions of the participants’ hands in response to an early versus late switch from a square to a rectangle target with rightward perturbation (Figure 3B, red and blue, respectively), a significant difference was observed (Kolmogorov-Smirnov test, p < 0.005, K = 0.4435). Differences in end-point distribution were also observed when the target goal was changed from a rectangle to a square (Figure 3E, red and blue, respectively) (Kolmogorov-Smirnov test, p < 0.005, K = 0.3780).

Similar observations characterized the responses to the target switches with leftward perturbations (Figure 3, C and F). For example, differences in the x-axis position of the participants’ hands at the end of the trials between switching conditions and the non-switching condition were also observed with leftward perturbation loads and with target switches from square to rectangle and vice versa (seeFigure 3, C and F). Thus, corresponding changes in control policies were elicited by both changes in target shape (from square to rectangle and vice versa) and following early and late switch conditions. Based on these observations, we next focused on the EMG data to gain insight into the latency of the changes in neural control evoked by switching of the target.

### Muscular activity

Based on the behavioral analyses presented above, we hypothesized that evoked EMG responses to perturbation loads would increase or decrease when the change in goal target increased versus when the spatial constraints imposed by the goal target were relaxed, respectively. A key aspect was to assess when a change in activity could be observed. Therefore, we initially examined global changes in EMG responses consistent with the changes in end-point distributions based on the activity averaged across different time windows after introduction of a perturbation load. Then, we used a sliding window analysis to measure the latency at which a change in the goal altered the EMG responses to perturbation. For this purpose, we examined stretching induced by rightward and leftward perturbations, respectively.

In Figure 4A and Figure 4B, responses for PM and PD are shown for the non-switching conditions, respectively. The EMG responses to the perturbation loads were higher in both PM and PD when the participants were reaching for the square target goal rather than for the rectangular target goal. A mixed model analysis employing bins of 25-ms were used to characterize these differences (see Methods). For rightward perturbations, the first significant larger PM response associated with the square target was observed between 125 and 150 ms (t(657) = -2.1446, p = 0.0323, d=0.19) (Figure 4A) followed by strongly significant differences in the subsequent bins(all p < 0.0005). Thirteen of the 14 participants exhibited individual differences in trial distributions. The first significant larger PD response was observed between 100 and 125 ms (t(657) = -4.0612, p < 0.005, d=0.43) (Figure 4D), also followed by strongly significant differences(all p < 0.005). Eleven of the 14 participants exhibited individual differences in trial distributions. These differences are slightly later that those reported in previous work [(Cross K.P., 2019) (Nashed J.Y., 2012) (Lowrey C.R., 2017)], which is likely attributable to the spatial layout, joint configuration and perturbation magnitude. A visual inspection indicated that trend started in the long-latency epoch without reaching significance (50-100 ms).

**Figure 4.**
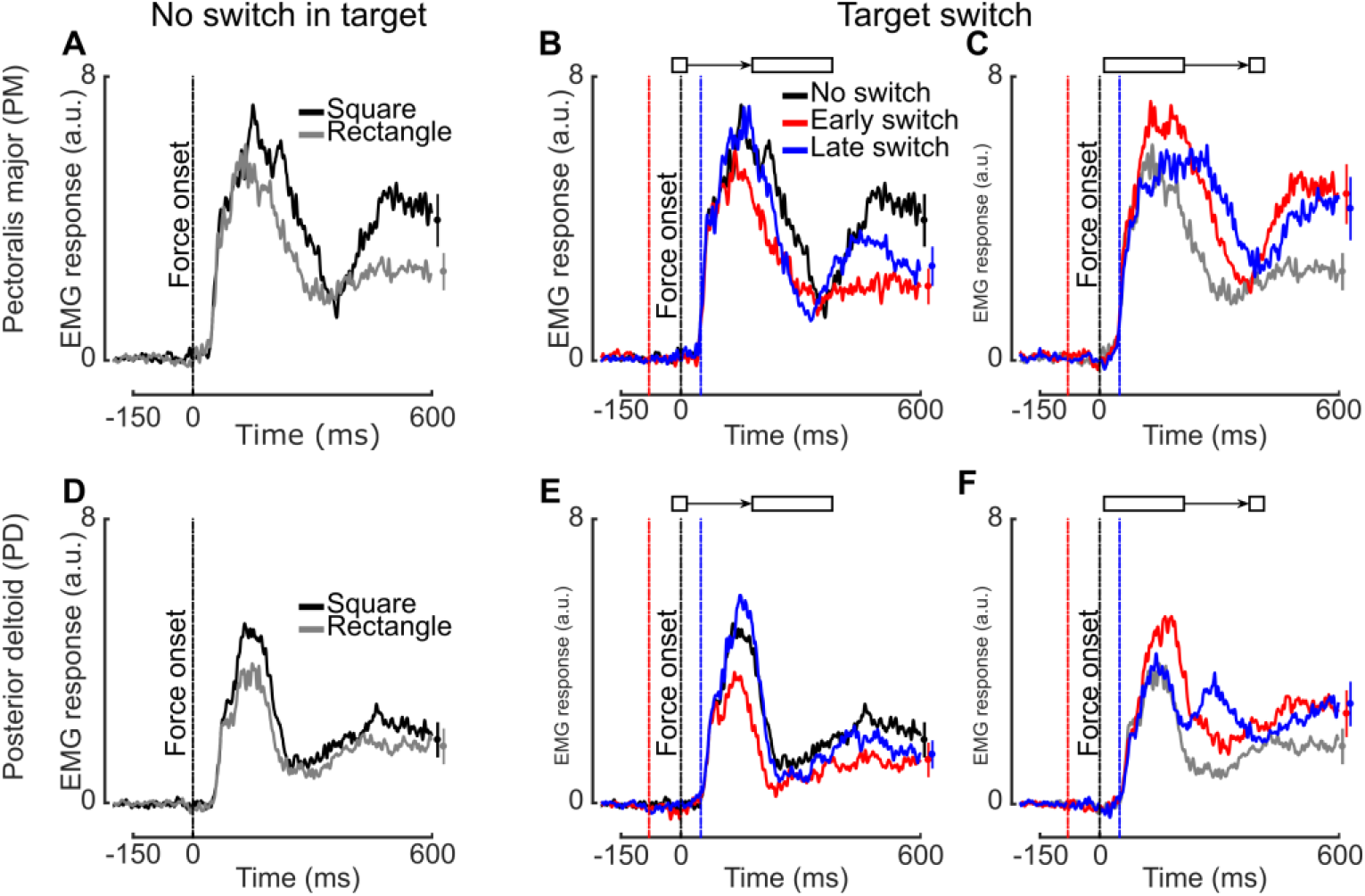
**A-F**. Mean baseline reduced EMG responses in the PM (**A**-**C**) or PD (**D**-**F**) of participants with a rightward (**A**-**C**) or leftward (**D**-**F**) perturbation load (indicated as a vertical dashed line) applied while reaching to a square (black), a rectangle (grey), or to a square switched to a rectangle (**B**,**E**) or vice versa (**C**,**F**), either early (red) or late (blue). For each plot, mean ± SEM values for the EMG responses 600 ms after the onset of force are shown at the far right.

Based on the observed EMG responses to perturbation loads for the square and rectangle targets separately, we hypothesized that a decrease in EMG response would be observed when the target was switched from square to rectangle, while an increase in response would be observed for a switch from rectangle to square. For PM and PD (Figure 4, B and E), the switch from square to rectangle elicited reduced EMG responses to perturbation loads. When both early (red) and late (blue) conditions were compared to the non-switching condition (black), it appeared that the control policy was adjusted during movement. For example, when an early switch was made, a decrease in EMG response was observed for all of the participants between the 125–150 ms bin (t(657) = -2.0591, p = 0.0399, d=0.26) and the 275–300 ms bin (p < 0.0005 for all of the intermediate bins). Five of the 14 participants exhibited individual differences in trial distributions. When a late switch was made, a decrease in the PM EMG response was observed across all of the participants between the 200–225 ms bin (t(657) = -2.5964, p = 0.0096, d=0.17) and the 275–300 ms bin (p < 0.005 for all of the intermediate bins). Seven of the 14 participants exhibited individual differences in trial distributions. Meanwhile, for PD, an early switch condition resulted in a decreased EMG response between the 100–125 ms bin (t(657) = -5.1381, p < 0.005, d=0.3) and the 250–275 ms bin (p < 0.005 for all of the intermediate bins). Seven of the 14 participants exhibited individual differences in trial distributions. However, under the late switch condition, no decrease in EMG response was observed for PD.

Conversely, when the target was switched from rectangle to square, PM and PD EMG responses to perturbation loads increased (Figure 4, C and F). The early (red) and late (blue) switch conditions were each compared to the non-switching condition (grey). For PM, the early switch condition resulted in an increase in EMG response from the 100–125 ms bin (t(657) = -2.4266, p = 0.0155, d=0.18) to the 275–300 ms bin (p < 0.0005 for all of the intermediate bins). Nine of the 14 participants exhibited individual differences in trial distributions. When a late target switch was introduced, a significant increase in EMG response was observed from the 150–175 ms bin (t(657) = - 2.6505, p = 0.0082, d=0.19) to the 275–300 ms bin (p < 0.005 for all intermediate bins). Eleven of the 14 participants exhibited individual differences in trial distributions. For PD, the early switch condition elicited an increase in EMG response from the 100–125 ms bin (t(657) = -3.2511, p = 0.0012, d=0.19) to the 275–300 ms bin (p < 0.005 for all intermediate bins). Ten of the 14 participants exhibited individual differences. Under the late switch condition, significant differences were observed from the 200–225 ms bin (t(657) = -3.7121, p = 0.0002, d=0.21) to the 275–300 ms bin (p < 0.005 for all intermediate bins) and 10 of the 14 participants exhibited individual differences in trial distributions.

The EMG and kinematics responses to perturbation loads both indicate that the participants adjusted their behavior during movement in response to a switch in target shape. These data also suggest that the time onset of those responses correlates with the onset of the target switch. Indeed, we observed that differences in both kinematics and EMG responses occurred earlier in the trials involving an early target switch than in those involving a late target switch. To better characterize the delay between the trigger of a target switch and the first differences in EMG responses, we subtracted the EMG responses obtained under non-switching conditions from those obtained under switching conditions. Thus, the remaining signals represented the responses to the target switch itself (Figure 5), and the onset of differences in the EMG responses to the target switch was found to depend on the condition. For example, the onset of differences in EMG response was associated with the first 25-ms bin in which the average EMG activity significantly differed from zero. Those bins are represented as shaded rectangles in Figure 5 for the different target switching conditions (e.g., early or late switching from square to rectangle or from rectangle to square) and for the two muscles (Figure 5, A and B for PM; Figure 5, C and D for PD). We also computed the individual mean values of the differences in EMG responses for the bins between 200 ms and 375 ms after a late onset of force (empty dots on the right side of each panel in Figure 5). When the square target was switched to the rectangle target, the mean values were negative (p < 0.005 in both muscles and for both early and late switch conditions). When the rectangle target was switched to the square target, the mean values were positive (p < 0.005 in both muscles and for both early and late switch conditions).

**Figure 5.**
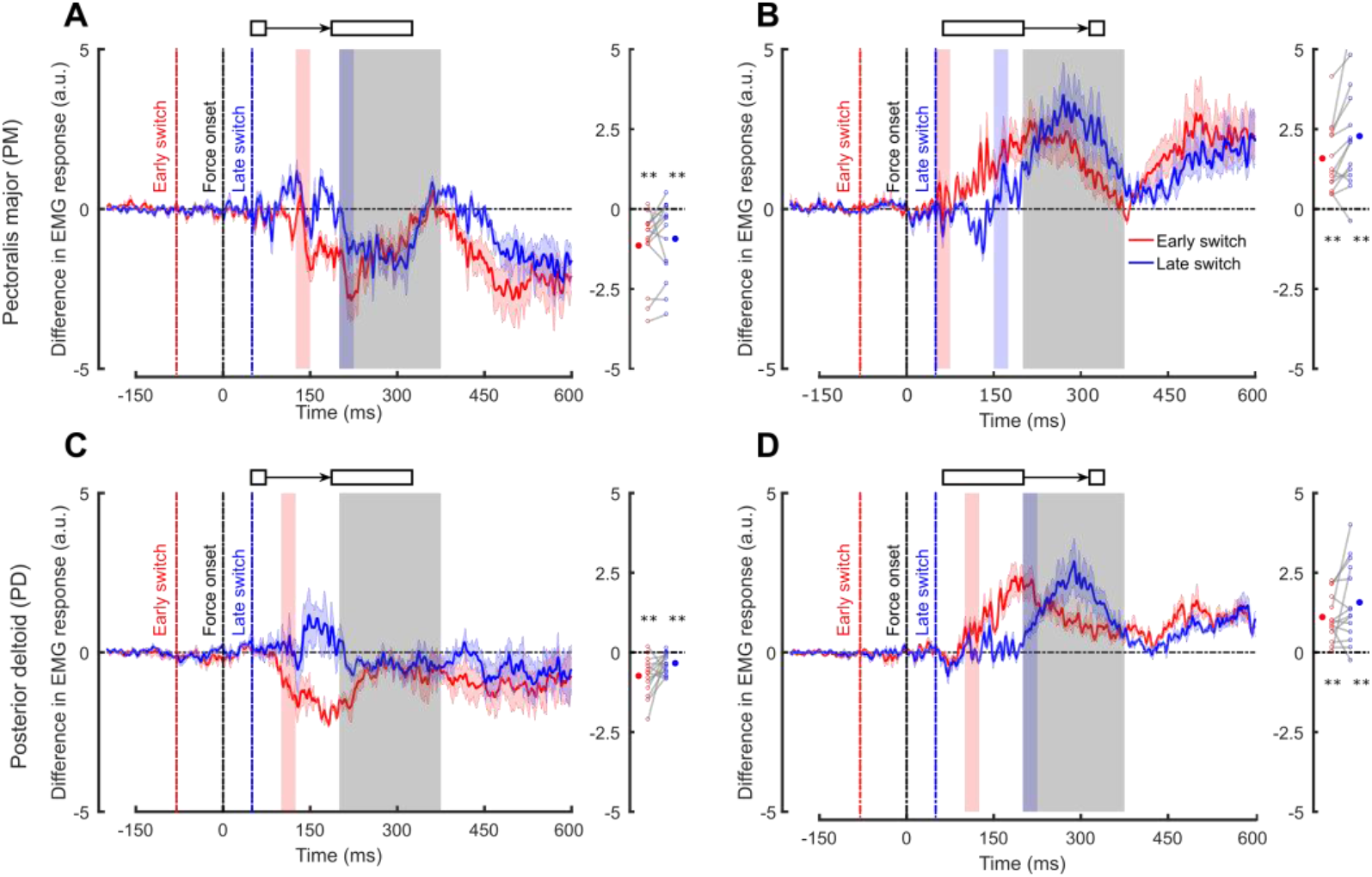
**A-D**. Mean and SEM differences in baseline reduced EMG responses in PM (**A**,**B**) or PD (**C**,**D**) while reaching to a square switched to a rectangle (**A**,**C**), or vice versa (**B**,**D**), either early (red) or late (blue). The onset of a rightward (**A**,**B**) or leftward (**C**,**D**) perturbation load is indicated with a black dashed line. The red and blue shaded rectangles in each panel represent the first significant bins for both conditions. At the far right of each panel, individual mean difference values in baseline reduced EMG are shown for the grey shaded rectangle for both early (red points) and late (blue points) conditions. The significance levels are **p < 0.005 and *p < 0.05 and these were assessed for all of the participants.

Thus, the results of Experiment 1 indicate that the participants adjusted their control policy during their movements when the goal target was switched from a square to a rectangle, or vice versa. Significant differences were also observed in both the kinematics and EMG responses in response to the two switching conditions. Delays between the trigger of the target shape switch and the first differences in EMG response spanned 150 ms to 200 ms. An analysis of the individual muscle sample data further indicated that changes occurred as early as 125–150 ms after the target switch. We believe that these timing for the first differences in EMG are meaningful number but we acknowledge that it should be taken with caution as there is some uncertainty linked to the bin width.

### Experiment 2

In order to demonstrate reproducibility and to test similar principles in a different context, we performed a second experiment which was designed to be a variant of Experiment 1. Briefly, we instructed participants to perform reaching movements to attain a square target in the presence of two virtual, rectangularly-shaped obstacles (see Methods). A lateral force was applied (as described in Experiment 1) to enhance possible changes in control which would be evoked by the appearance or disappearance of the obstacles. Changes in control policy were assessed based on movement kinematics and evoked EMG responses of the muscles stretched by the perturbation.

An analysis of the participants’ movement kinematics when the obstacle conditions remained unchanged revealed clear differences between the trials with and without obstacles. First, both the hand paths (Figure 6 A-F) and end-point hand distributions significantly differed when obstacle conditions were changed (appearing vs. disappearing, Peacock test). These kinematics results confirmed that the control policy of ongoing movements is adjusted depending on whether obstacles are present.

**Figure 6.**
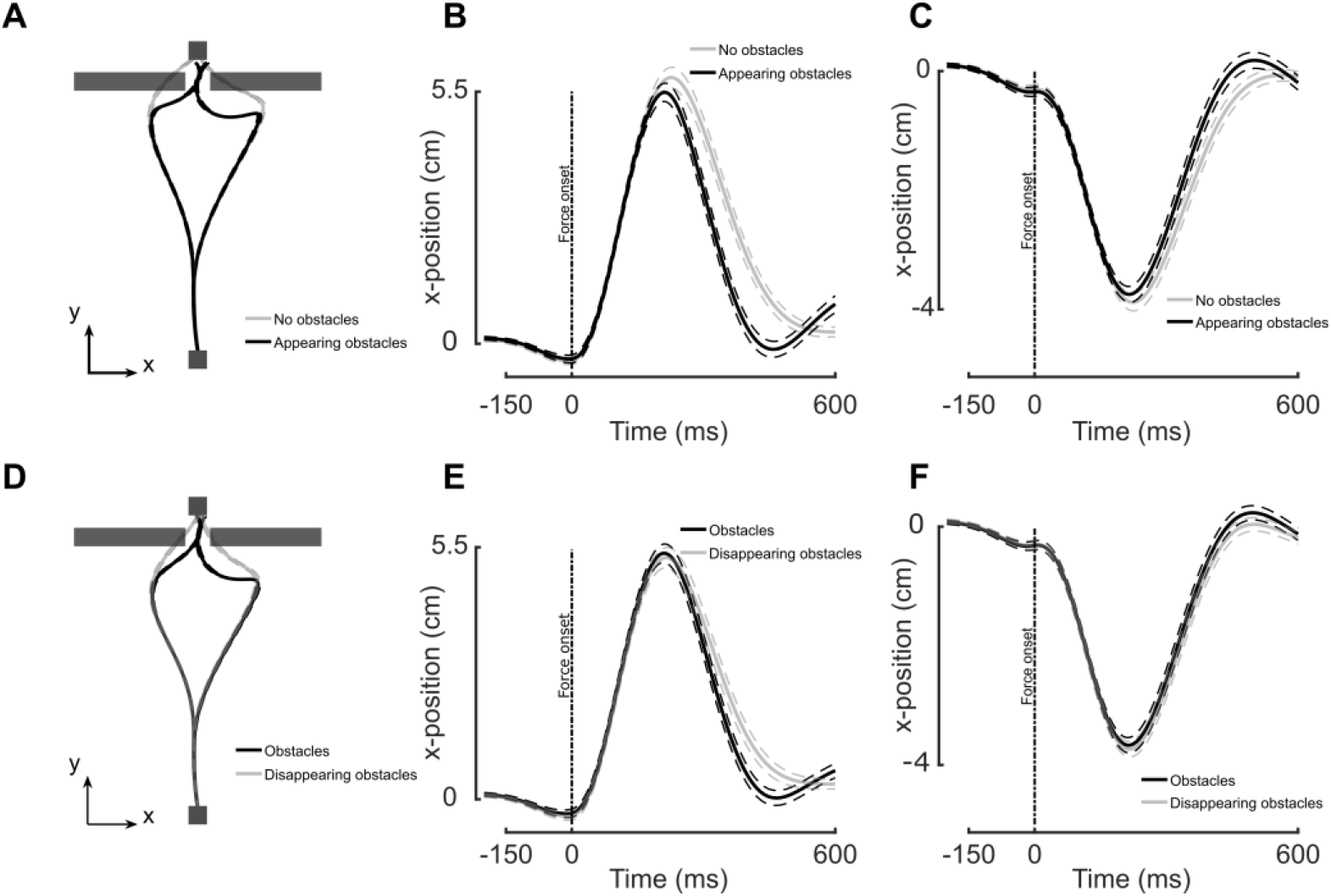
**A-D**. Mean (solid line) and SEM (dashed line) traces of participants’ hand paths (**A**,**D**) and x-axis positions (**B**,**C**,**E**,**F**) with respect to time in reaching for a target with a perturbation load applied in the presence (black) or absence (grey) of obstacles as indicated

Similar to Experiment 1, we also examined evoked EMG responses of the PM and PD muscles stretched by perturbations. Greater activity was detected in the presence of the obstacles (black) than in the absence of the obstacles (grey). Linear mixed model analysis revealed the first significant difference between conditions in the time bin 50-75 ms (p<0.005) for the Pectoralis major. Figure 7A and 7D present the responses for PM and PD, respectively. Differences were also observed in the evoked EMG responses to the perturbation loads when the obstacles appeared versus disappeared. For example, the first increase in EMG response for PM was observed when obstacles appeared during the 100–125 ms bin (t(798) = 2.4816, p = 0.0133, d=0.11) and for PD during the 150–175 ms bin (t(798) = 2.0009, p = 0.0497,d=0.10). Then, when the obstacles disappeared, the first decrease in EMG response in PM was observed during the 150–175 ms bin (t(798) = -2.3674, p = 0.0181, d=0.11) and in PD during the 150–175 ms bin (t(798) = 2.1805, p = 0.0295, d=0.11). These results in the presence and absence of obstacles are shown in Figure 7B, E, C and F respectively. We further observed, similar to Experiment 1, that individual differences in trial distributions were detected for eight, seven, six, and seven participants, respectively.

**Figure 7.**
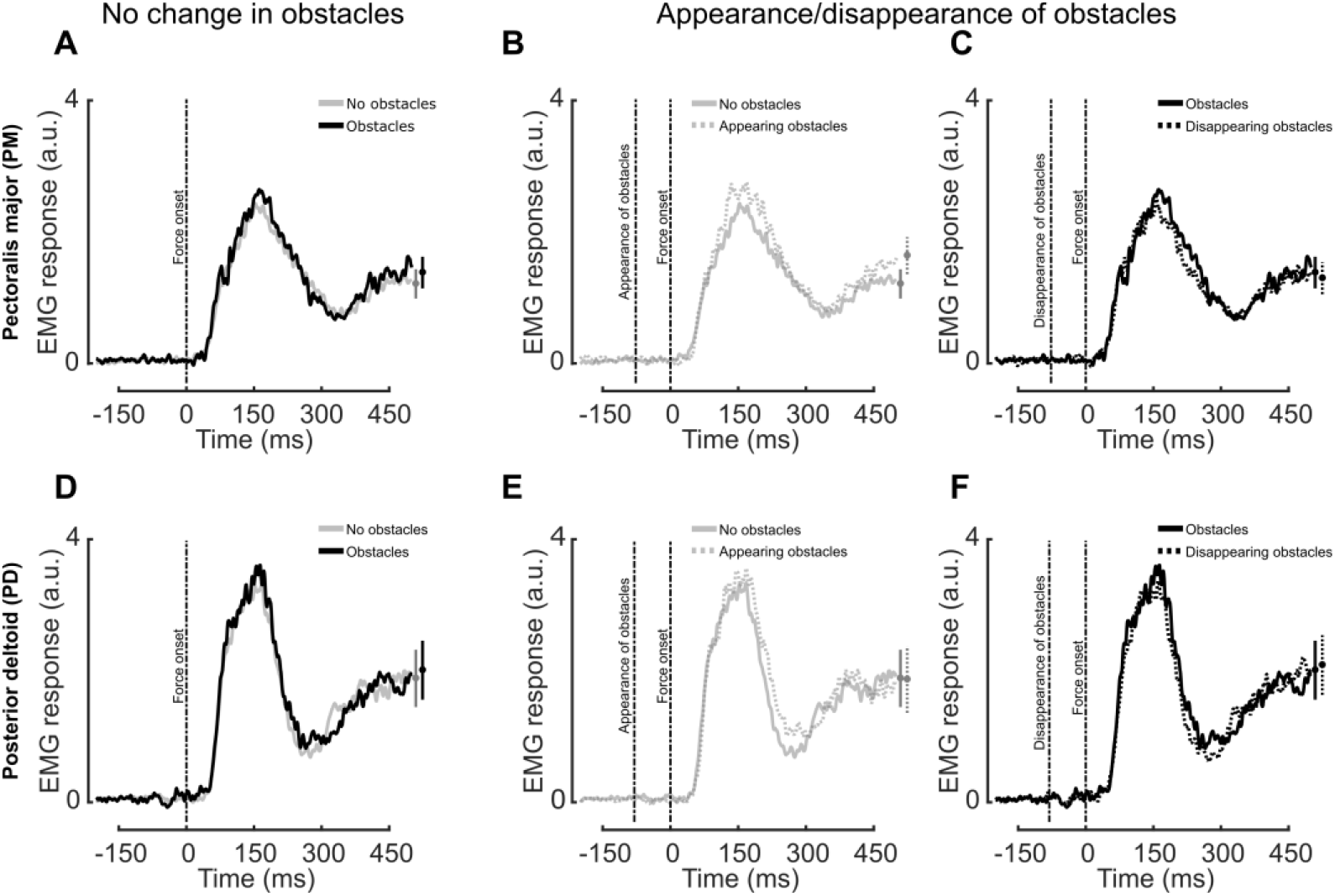
**A-F**. Mean baseline reduced EMG response traces measured in the presence or absence of obstacles as indicated for PM (**A**-**C**) or PD (**D**-**F**) with a rightward (**A**-**C**) or leftward (**D**-**F**) perturbation load applied. Vertical dashed lines are used to indicate the time of force onset and the appearance or disappearance of obstacles as labelled. Mean ± SEM values are shown at the far right of each plot.

Also similar to Experiment 1, data for the kinematics and EMG responses to perturbation loads indicate that participants adjusted their policy during movement in response to changes in context (e.g., the appearance/disappearance of obstacles). To obtain an approximation of the delay between the appearance or disappearance of obstacles and the differences in EMG responses, we subtracted the EMG responses obtained when there was no change in the obstacles from the EMG responses obtained when the conditions were changed. Specifically, for the trials involving the appearance of obstacles, the no obstacles condition was subtracted; for the trials where the obstacles disappeared, the obstacles condition was subtracted. The resulting signals corresponded to the impact of rapid changes in context (Figure 8). Differences in EMG response occurred within the first 25 ms in which a significantly non-zero average EMG activity was detected. The relevant bins are represented as shaded rectangles in Figure 8. Observed delays between the onset of the appearance or disappearance of obstacles and the first differences in EMG responses spanned an interval of 180 ms to 250 ms.

**Figure 8.**
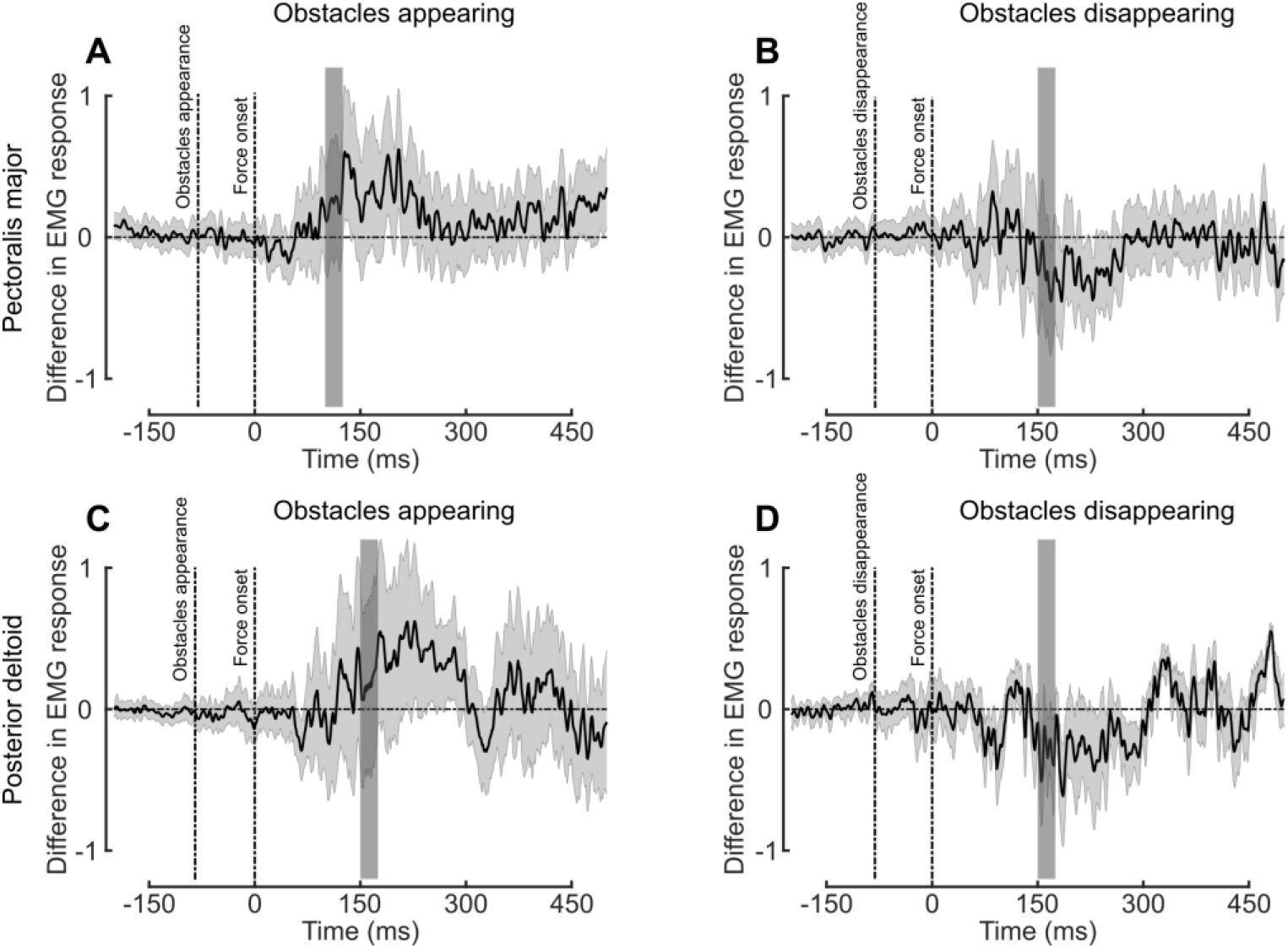
**A-D**. Mean and SEM differences in baseline reduced EMG responses in PM (**A**,**B**) or PD (**C**,**D**) due to an appearance (**A**,**C**) or disappearance (**B**,**D**) of obstacles with rightward (**A**,**B**) or leftward (**C**,**D**) perturbation loads. The shaded rectangles in each panel represent the first time bin in which the response significantly differs from zero.

### Control experiment

In order to verify that the change in behaviour observed in Experiment 1 could be elicited without mechanical perturbations (see Methods), we performed a third (control) experiment that was a variant of Experiment 1, in which there was no mechanical perturbation. Participants performed reaching movements to a goal target that could either be a small square or a large rectangle. During movement, the target shape could switch from square to rectangle and vice-versa. The occurrence of a change in behavior induced by this switch in target shape was assessed based on hand kinematics. Figure 9A and 9B represent the hand paths of a representative subject toward a square that turned into a rectangle and a rectangle that turned into a square respectively.

**Figure 9.**
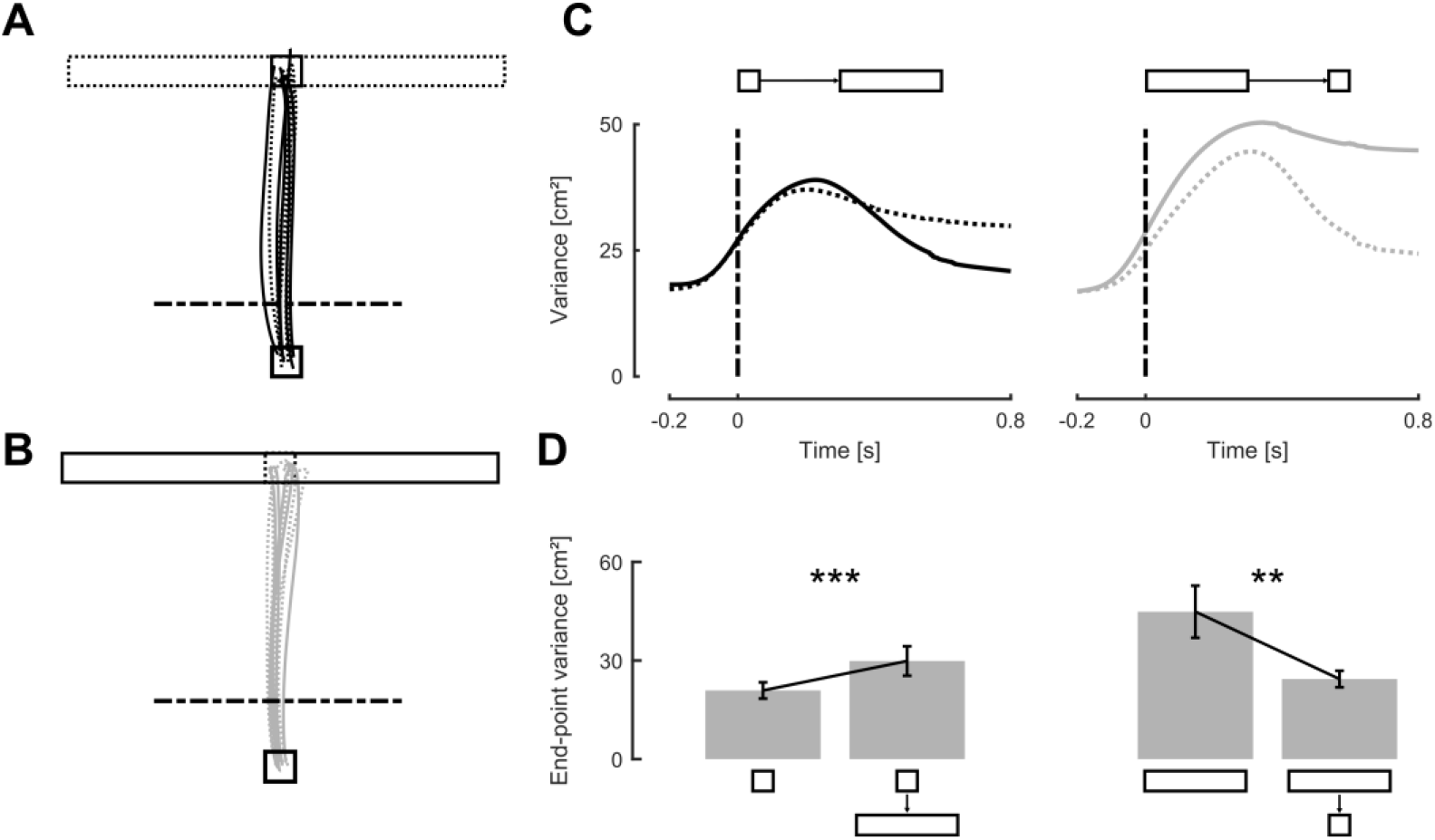
**A-B**. Hand paths of a representative participant toward a square (**A**) or rectangle (**B**) target, full lines correspond to the non switching condition and dashed lines correspond to the switching condition. Horizontal dashed lines represents the position of the onset of the visual perturbation. **C**. Evolution of the hand path variance along the x-axis for the unperturbed condition (full line) and the switching condition (dashed) for initial square (Left) and rectangle (Right) targets. **D**. Mean and SEM of the end-point variance of participants’ hand position along the x-axis. The left barplot represents the unperturbed condition and the right bar plot the perturbed position.

The end-point distribution of participants’ hand for movements without target switch was different for square and rectangular targets as expected. The medians of these end-point distributions for rectangle and square targets were not significantly different (Wilcoxon signed rank test, z = 1.86, ranksum = 2.38∗ 10^5^, p=0.0618). However, we observed a wider distribution for rectangle (−0.7 ± 5.3 cm) than for square targets (−0.03 ± 0.35 cm) across subjects (two-samples Kolmogorov-Smirnov test, p<0.005, K=0.1125). One-tailed paired t-test performed on individual variances confirmed this finding (p<0.005, t = -4.02, d=1.18). Strikingly, target switches online also elicited changes in end-point variance (Figure 9C). Indeed, we observed an increase (one-tailed paired t-test: p<0.005, t=-4.10, d=- 0.82) or a decrease (one-tailed paired t-test: p<0.005, t= 3.46, d=1.06) in end-point variance when the target switched from square to rectangle or from rectangle to square respectively (Figure 9D).

To conclude, online modification of the goal target or context elicited online changes in control, impacting corrections for lateral deviations due to natural variability (control Experiment) as well as feedback corrections to external loads applied during movements (Experiments 1 and 2).

## Discussion

We conducted three experiments to determine whether and how quickly the control strategy used by humans during reaching can be adjusted online when the structure of the target goal, or the environmental context, unexpectedly changes. In Experiment 1, we changed the structure of the goal target from a small square to a large rectangle and vice versa. In Experiment 2, obstacles located on both sides of the straight path to the target suddenly appeared or disappeared during movement. In both experiments, we found that the control strategies used by participants adjusted online following the changes in target shape or context. In Experiment 1, the evoked EMG responses to mechanical perturbations adjusted in as little as 150 ms following the change in target structure. Similarly, in Experiment 2, adjustments of the responses to mechanical perturbations were observed 180 ms after the appearance or disappearance of obstacles. Finally, in Experiment 3, we observed a change in movement execution in response to the same target switch as in Experiment 1, even in the absence of mechanical perturbation. This shows that the evoked modulation of feedback gains is not only linked to feedback responses but instead it also reflects a more general adjustment of control expressed in both perturbed and unperturbed movements.

This interpretation is in line with the theory of optimal feedback control which suggests that the control strategy used to perform movement and to correct for errors is tuned to the task goal and to its environmental context (Liu and Todorov 2007; Scott 2004; Todorov and Jordan 2002).

Mathematically, the tuning of the control policy reflects the minimization of a cost-function, which captures the intended behavioral performance by weighting motor cost and penalties on motor errors (e.g. position, velocity, and force). In order to model the structure of the goal target in our experiments, a selective penalty on the x-coordinate for the square versus rectangle target could have easily been introduced. However, this modeling approach would be more complicated for the obstacle conditions since non-quadratic penalties would be involved but clearly context-dependent changes in control strategy are expected.

In this framework, our experiments indirectly probed the two terms typically used in standard OFC models: a term related to behavioral performance that penalises error in the state and a term related to motor cost that penalises control commands. On the one hand, a switch from a rectangle to a square, or an appearance of obstacles, clearly alters the penalty related to the state. On the other hand, when the target switched from a square to a rectangle, or when the obstacles disappear, adjustments of control commands were not mandatory. The visual perturbation only modified the penalty on behavioral performance and left the penalty on motor cost unaffected, but the relative weight of motor cost likely promoted an adjustment of the control policy when the task became easier, as following a switch from a square to a rectangle, or when the obstacles disappeared. The net result is a decrease in EMG-evoked responses. This adjustment is observed with the rightward perturbation for both early and late target switches (Figure 4B, red and blue traces, respectively), yet only for the early target switch with leftward perturbation (Figure 4E, red trace). Moreover, even though the penalty regarding behavioral performance decreased, participants did not have to adjust their control policy to succeed in the trial. The fact that those adjustments are not mandatory, may explain why the decrease observed was not strong in all cases at short latencies, although an overall decrease later in the trial was always observed (right part of Figures 5A and 5C).

Many studies have investigated the impact of target jumps (Georgopoulos et al. 1981; Goodale et al. 1986; Gritsenko et al. 2009; Gritsenko and Kalaska 2010; Mutha et al. 2008; Pélisson et al. 1986; Smeets and Brenner 2003) and cursor jumps (Franklin and Wolpert 2008; Sarlegna et al. 2003) on control of reaching movements. In a recent study, these two types of visual perturbations were combined to test whether a motor system was able to update its control strategy during movement (Dimitriou et al. 2013). However, it was recently demonstrated that neither target jumps nor cursor jumps necessarily imply a modification of the objective function, and therefore, they do not necessarily involve an update in control strategy (Li et al. 2018; Qian et al. 2013). A key aspect of the present work was that both a specific change in target structure and a change in environmental context were examined. Based on previous empirical and theoretical studies (Lowrey et al. 2017; Nashed et al. 2012), we included both online modification of target structure, as well as online appearance or disappearance of obstacles. Unlike a target jump, these two changes in task demand cannot be captured by a change in the state of the target or cursor with a single closed-loop controller. We also sought to measure the latency at which the control policy in the motor system was updated. In the first two experiments, the average latencies observed in the EMG responses were surprisingly short: 150 ms in Experiment 1 and 180 ms in Experiment 2. In comparison, visuomotor corrections by muscles in a target jump paradigm emerge within ∼100 ms after the target jump (Day and Lyon 2000; Franklin and Wolpert 2008; Knill et al. 2011).

Interestingly, the difference of 30ms observed across Exp. 1 and 2 may be due to differences in the tasks performed. While the first change only affects the task goal, the second change affects the environmental context. A recent study also reported an additional 30 ms latency in feedback responses to cursor jumps when the response was influenced by the presence of obstacles in the context of motor responses to visual perturbations (Cross et al. 2019). Indeed, Cross and colleagues have shown that the motor response to a visual perturbation includes two distinct phases. The first phase starts after 90 ms and is sensitive to goal redundancy. Meanwhile, the second phase starts after 120 ms and it is sensitive to environmental factors. Similarly, in the present study, a change in goal and a change in context resulted in a 30 ms difference in latency. Thus, it is conceivable that 30 ms is a common additional processing time for tasks involving environmental obstacles. Considering a fixed visuomotor delay of ∼100 ms (Franklin and Wolpert 2008; Saunders and Knill 2004; Smeets and Brenner 2003)], and 30 ms across experiments linked to a specific processing of obstacles, the measured latencies of target-dependent responses suggest that a change in control policy requires 50 ms.

In order to set constraints on the online processes that adjust control policy, we compared the latencies observed in the present study with the visuomotor delays previously measured with target or cursor jumps and also with reported reaction times required to initiate a movement. The latencies we observed were longer than visuomotor delays associated with target or cursor jumps (∼100 ms), which suggests that a more demanding process than the one involved in target jumps is used. On the other hand, the latencies we measured were shorter than the reaction times characterized for initiating a movement (Georgopoulos et al. 1981). The latter result suggests that the control policy is not re-computed from scratch. However, this assessment has to be taken with cautious since Haith and colleagues (Haith et al. 2016) showed that these reactions time not only reflect control policy selection and that movement preparation can occur in as little as ∼130 ms. It is also important to stress out that the reaction times we used for these comparisons were mostly measured for transitions from postural control to reaching movements. In a previous study (Orban de Xivry et al. 2017), reaction times were reduced if movement planning and movement execution overlapped. This result is consistent with the present experiments since our participants had already started their movement when they had to adjust their control policy.

Two different kinds of adjustments in control policy could occur during the movement in our experiments. The first is re-computation of a new control policy for the new task, corresponding to the framework of model predictive control (Lee 2011). Many recent theories of motor control consider that the control policy is computed before movement is initiated (Harris and Wolpert 1998; Scott 2004; Todorov and Jordan 2002; Wong et al. 2015). It has recently been suggested that specification of this control policy may overlap with movement execution (Orban de Xivry et al. 2017), which could support hypothesis about the process that perform the adjustment in control policy. The second process which could produce an online adjustment of control policy is a switch between different control policies stored in the brain. It has already been suggested that the central nervous system is able to change online the target goal of a movement if a perturbation interferes sufficiently with the task (Nashed et al. 2014). Other recent studies have suggested that multiple motor plans may be handled in parallel by the central nervous system (Gallivan et al. 2016b, 2016a). The present data do not allow us to determine whether the observed adjustments corresponded to a switching or re-computation. Thus, further studies are needed to further elucidate the details of this process.

Interestingly, when the goal target was switched from a rectangle to a square in Experiment 1, we observed a shorter latency of the EMG-evoked motor response for the late switching condition that for the early one. This difference may reflect the participants’ urgency in correcting their movement toward the new goal target. Indeed, long-latency responses and early voluntary epochs of responses to perturbations have been shown to scale with task-related urgency (Crevecoeur et al. 2013). In the present study, the participants’ urgency to correct mechanical perturbations was larger when the switch from a rectangle target to a square target was made late in the trial. This result could explain the smaller latency of the response to the perturbation load. Another factor which may have potentially influenced these differences in latencies is the temporal evolution of feedback gains during reaching. It has been shown both experimentally and theoretically that feedback gains follow a skewed bell shape curve over time (Dimitriou et al. 2013; Liu and Todorov 2007). As a result, different responses to the same perturbation are applied at distinct points in time. In our experiment, the same differences were observed when the target was switched from a square to a rectangle.

Understanding the neural basis of the change in movement control remains an open question, however our developments provide a measurement of the latency and sets constraints on candidate underlying circuits. On the one hand, both spinal and supraspinal circuits are recruited during movement and following disturbances, but when the target did not change, the task-dependency emerged in the long-latency and voluntary epochs, consistent with previous work (Lowrey et al. 2017) and suggesting a contribution of cortical origin (Matthews 1991; Pruszynski et al. 2011; Scott 2004). On the other hand, the main timing constraint in the condition where the target switched was due to the processing of the visual system. Rapid reflexive responses to visual perturbations are typically evoked in a little less than 100 ms (Day and Lyon 2000; Franklin and Wolpert 2008; Knill et al. 2011), but we observed longer latencies for task-related changes likely due to more demanding processing associated with a change in control strategy. In fact, our measure of the latency associated with changes in the shape of the target involves both visual and somatosensory systems, thus it is reasonable to speculate that associative regions in parietal cortex play a central role. With regards to timing, this suggestion is consistent with a cortical pathway collecting sensory signals from V1 and S1 to modulate the generation of motor commands through M1 (Omrani et al. 2016).

To conclude, we have demonstrated that humans are able to quickly update their control policy when a task goal or its environment changes during reaching movements, and that these changes characterize unperturbed movements as well as feedback responses to mechanical loads applied to their limb. We observed adjustment delays which were slightly longer than the visuomotor delays observed in a target jump paradigm, which suggests that a more demanding neural operation is needed to the change in control policy. Our estimate of the latency associated with this process is ∼30 ms, therefore allowing fast motor adjustments suitable for the execution of ongoing movements in a dynamic context. These valuable insights provide direction for further studies of movement planning and how it interacts with movement execution dynamically.

